# Platinum resistance induces diverse evolutionary trajectories in high grade serous ovarian cancer

**DOI:** 10.1101/2020.07.23.200378

**Authors:** J.I. Hoare, H. Hockings, J. Saxena, V.L. Silva, E. Maniati, H.B. Mirza, W. Huang, G.E. Wood, F. Nicolini, T.A. Graham, I.A. McNeish, M. Lockley

## Abstract

Resistance to therapy is an enduring challenge in cancer care. Here we interrogate this critical unmet need using high grade serous ovarian cancer (HGSC) as a disease model. We have generated a unique panel of platinum-resistant HGSC models and shown that they share multiple transcriptomic features with relapsed human HGSC. Moreover, they evolve diverse *in vivo* phenotypes reflecting the human disease. We previously characterised copy number signatures in HGSC that correlate with patient survival and now provide the first evidence that these signatures undergo recurrent alterations during platinum therapy. Furthermore, specific, resistance-associated signature change is associated with functionally relevant gene expression differences. For example, reduced signature 3 (BRCA1/2-related homologous recombination deficiency) is associated with increased expression of homologous recombination repair genes (Rad51C, Rad51D, BRCA1) and DNA recombination pathway enrichment. Our mechanistic examination therefore provides new and clinically relevant insights into the genomic evolution of platinum-resistant cancers.

## Introduction

Epithelial ovarian cancer is a heterogeneous disease, including five different pathological subtypes, the most common of which is High Grade Serous Cancer (HGSC) [1]. Standard treatment to date has been combination surgery and platinum-based chemotherapy. Despite this aggressive approach, 85% of patients will relapse and respond less well to successive platinum-containing chemotherapy regimens [2]. This inevitable development of chemotherapy resistance is thus an unmet clinical need and five year survival in advanced disease is static at approximately 15% [3].

HGSC is characterised by mutations in TP53 [4]. It has been divided into four molecular groups based on gene expression profiles; immunoreactive (C2), proliferative (C5), differentiated (C4) and mesenchymal (C1) [5]. The Cancer Genome Atlas Research Network (TCGA), confirmed near ubiquitous TP53 mutation (96%) as well as increased copy number variation. Germline loss of function or methylation was commonly observed in BRCA1 (9%) and BRCA2 (8%), while defective homologous recombination repair pathways occurred in 49% of cases [6]. These genetic discoveries have been instrumental in the clinical implementation of PARP inhibitor therapy in HGSC, which has led to dramatic improvements in progression-free survival and changed the treatment landscape in platinum-sensitive disease [7-12].

Despite these notable advances in the genetic and molecular characterisation of HGSC, new treatments have not improved outcomes for patients with platinum-resistant cancer. To address this, we and others have interrogated the genomic features of HGSC, including in relapsed disease [13, 14]. The clinical phenotypes observed in platinum-resistant HGSC vary widely so that some women may experience relapse with widespread metastasis and recurrent ascites, while others experience oligometastatic recurrence, usually with a more indolent disease course. Although we have recently defined copy number signatures and shown that they correlate with patient survival [14], the specific role of these genetic changes in directing the evolution of diverse clinical phenotypes has not previously been interrogated.

Pre-clinical research relies on cell lines but the extent to which they reflect features of relapsed, human HGSC is largely unknown and the routine creation of permanent cell lines from human HGSC has proved to be extremely challenging. Although the advent of novel culture media has improved success rates [15], when compared to established cell lines, primary cells are limited by ethical difficulties in access for international researchers and a lack of publicly available annotated data [16]. Most laboratory research in ovarian and other cancers has relied heavily on commercially available cell lines. Domcke *et al* [17] compared the genomes of 47 ovarian cancer cell lines to the TCGA dataset and demonstrated that the most commonly used cells, including A2780 and SKOV3, represent human HGSC poorly. Cell lines that Domcke *et al* defined as “likely high grade serous” included OVCAR4, Cov318 and Ovsaho and subsequent work has demonstrated their utility as laboratory research tools [16, 18].

Here we use these same cell lines [17] to create a novel panel of platinum-resistant *in vitro* and *in vivo* HGSC models that recapitulate the genetic and clinical features of the human disease. By exploiting this unique resource, we interrogate for the first time how drug pressure influences multiple genomic features, including our novel copy number signatures [14], and show how these changes relate to the evolution of distinct clinical phenotypes.

## Methods

### Cell culture and chemotherapy-resistant cell lines

Human HGSC cell lines OVCAR4, Cov318 and Ovsaho were obtained from Prof Fran Balkwill (Barts Cancer Institute, UK) and grown in DMEM (OVCAR4 and Cov318) or RPMI (Ovsaho) media containing 10% FBS and 1% penicillin/streptomycin. Platinum-resistant HGSC lines were generated by culturing cells in increasing concentrations of cisplatin or carboplatin at each passage of confluent plates until a fold increase in IC_50_ of 2-10 was achieved to mirror published estimations of the resistance observed in human patients [19]. All cell lines underwent 16 locus STR verification (DNA Diagnostics Centre, London, UK: June 2015-February 2016 and European Collection of Authenticated Cell Lines August 2019) and routine mycoplasma testing. Spheroids were created using ultra-low attachment flasks (Corning) and media supplemented with 1% Insulin-Transferrin-Selenium (ITS) (ThermoFisher), 2% B-27 supplement (Life Tech), 20ng/ml EGF (Fisher Scientific) and 20ng/ml FGF (Fisher Scientific).

### Cell viability

Cell viability was assessed either with MTT (Thiazolyl Blue Tetrazolium Bromide, Alfa Aesar) or CTG, (CellTiter-Glo^®^, Promega). For MTT, cells were plated in triplicate in 24-well plates and treated the following day with cisplatin, carboplatin or vehicle. MTT (5mg/ml) was added 72 hours post-chemotherapy, and after two hours of drug incubation, crystals were dissolved in DMSO and absorbance measured using a Perkin-Elmer plate reader. For CTG, cells were again treated with chemotherapy or control and CTG reagent was added 72 hours post-chemotherapy. Light emission was quantified using a Perkin-Elmer plate reader. In all cases, dose-response curves were generated and IC_50_ calculated using GraphPad Prism v7.04.

### IncuCyte^®^

To measure proliferation, cells were plated in 96-well plates and incubated in the IncuCyte^®^ZOOM machine. Four images per well were recorded every four hours for five days and a confluence mask was created using IncuCyte^®^ ZOOM 2016B software. To assess cell migration, 2µg/ml MitomycinC was added for two hours before creating homogeneous, 700-800 μm-wide scratch wounds in confluent wells using the IncuCyte WoundMaker^®^. Wound density was measured over time using the IncuCyte^®^ live cell analysis imaging system.

### Organotypic cultures

Optically clear 0.4 µm pore transwells (Corning) were coated with 1% collagen in filtered PBS and incubated at 37°C for 1 hour. 3:1 Matrigel/collagen gels were mixed in 10x DMEM and 10% FBS at neutral pH. 100µl of gel mixture was added to each collagen-coated transwell and left to set for two hours at room temperature. Primary human fibroblasts were isolated from the metastatic omentum of women with HGSC in collaboration with Prof Fran Balkwill (Barts Cancer Institute, UK). Samples were collected with informed consent, via the Barts Gynae Tissue Bank (Research Ethics Committee: 15/EE/01/51). Briefly, omental tissue was submerged in 5X trypsin for 20 minutes before being diced, re-suspended in 0.5mg/ml collagenase (ThermoFisher) and digested with shaking (55rpm for 75 mins at 37°C). Digested tissue was filtered through 250 μl tissue strainers. Adipocytes were collected, re-suspended in 10ml DMEM+10% FBS and transferred into a flask as the fibroblast fraction. 1:2 mixtures of HGSC cell lines : human primary fibroblasts (3.3×10^4^ cells total in 200µl) were added to each pre-prepared gel. On day nine, gels were fixed in formalin prior to paraffin-embedding and staining with haematoxylin and eosin (H+E) and immunohistochemistry (IHC) for p53 (Dako IS616). Stained sections were imaged with a Pannoramic 250 scanner (3DHISTECH Ltd) and images analysed using Image J^®^ software. Invasion index was calculated as: (cell number) x (area) x (maximum depth) [20].

### Luminescent and fluorescent reporters

OVCAR4 cells were treated with lentiviral particles (MOI 10) containing dual constructs of green fluorescent protein (GFP) and Firefly luciferase (CMV-Luciferase (firefly)-2A-GFP (Puro)) to create OVCAR4-Luc. Ov4Cis and Ov4Carbo cells were transfected with dual constructs of red fluorescent protein (RFP) and Firefly luciferase (CMV-Luciferase (firefly)-2A-RFP (Puro)) (AMS Biotechnology^®^) to create Ov4Cis-Luc and Ov4Carbo-Luc respectively. Transfected cells were cultured in puromycin-containing media (0.9 µg/mL).

### Animal studies

Experiments were conducted under UK government Home Office license (P1EE3ECB4). Cells were inoculated by intraperitoneal (IP) injection (5×10^6^ cells/200µl sterile PBS) or by subcutaneous (SC) injection (5×10^6^ cells/200µl) into both flanks of female CD1*nu*/*nu* mice (Charles River Laboratories). Subcutaneous tumour growth was monitored with callipers. To measure light output, animals received D-Luciferin monopotassium salt (ThermoFisher) 3.7mg/200µl PBS IP and light emission was recorded using an IVIS^®^ Spectrum (PerkinElmer) and analysed using Living Image^®^ v7.4.2. Mice were assessed for weight, general health, and accumulation of ascites and were killed according to UK Home Office guidelines. At necropsy, murine tissue was fixed in 10% formaldehyde and paraffin-embedded. 4µm sections were stained with H+E and by IHC for p53 (DAKO IS616), Pax8 (Abcam ab181054) and Ki67 (Abcam ab15580). Slides were examined by a specialist Gynae-oncology histopathologist (JMcD).

### Production of *ex-vivo* cell lines

Murine IP tumours were minced with a scalpel in ice-cold PBS and enzymatic digestion was performed using collagenase type I (Fisher: 1mg/ml in 5% FCS/PBS). Tissues were incubated at 37°C with shaking (90 rpm for over 1 hour) then strained through a 70µm filter, washed and plated in DMEM containing 10% FCS and 1% penicillin/streptomycin.

### RNA Sequencing and bioinformatics analysis

RNA was extracted using the Qiagen RNeasy kit according to the manufacturer’s instructions. RNA was quantified by NanoDrop™ spectrophotometer (Thermo) and quality determined on the Agilent Bioanalyzer 2100 using RNA Nano Chips. Library prep and RNA sequencing were carried out by the Oxford Genomics Centre (Wellcome Centre for Human Genetics, Oxford) using PolyA capture. Sequencing was performed to ∼32x mean depth on the Illumina HiSeq4000 platform, strand-specific, generating 75 bp paired-end reads. RNASeq samples were mapped to the human genome (hg19, Genome Reference Consortium GRCh37) in strand-specific mode as part of the Wellcome Trust Centre pipeline. Number of reads aligned to the exonic region of each gene were counted using htseq-count based on the Ensembl annotation. Only genes that achieved at least one read count per million reads (cpm) in at least twenty-five percent of the samples were kept. Conditional quantile normalization [21] was performed accounting for gene length and GC content and a log_2_-transformed RPKM expression matrix was generated. Differential expression analysis was performed in EdgeR using the ‘limma’ R package [22]. To identify gene clusters segregating multiple sample classes we used the function sam from R package siggenes. Gene-set enrichment analysis (GSEA) was performed using the GSEA software [23] to identify dysregulated canonical pathways curated in the Molecular Signatures Database (MSigDB-C2-CP v6.2) (FDR q < 0.05). RNA-Seq data have been deposited in Gene Expression Omnibus (GEO) under the accession number GSE141630.

The ICGC_OV read counts were extracted from the exp_seq.OV-AU.tsv.gz file in the ICGC data repository Release 20 [24]. Only genes that achieved at least one read count in at least ten samples were selected, producing 18,010 filtered genes in total and voom normalization was applied. Differential expression analysis was performed in EdgeR comparing platinum resistant plus refractory (N = 49) versus sensitive (N = 31) donor samples. All graphics and statistical analyses were performed in the statistical programming language R (version 3.1.3).

### DNA extraction and deep whole genome sequencing (dWGS)

DNA was extracted from cultured cells using a QIamp DNA Blood Mini Kit (Qiagen) according to the manufacturer’s instructions. DNA quantity was evaluated using the Qubit 4 Fluorometer (Invitrogen™) then sonicated to fragments of 300-400bp using the M220 focused ultrasonicator (Covaris). Sequencing libraries were prepared using the NEBNext Ultra kit (New England Biolabs), according to the manufacturer’s instructions using 300ng fragmented DNA input for each sample. Libraries were sequenced to a target coverage of 50x on Illumina’s HiSeq X Ten platform (150bp paired end reads).

Platypus was used for variant calling of point mutations by comparison to the human reference genome (hg19). Mutations were considered if the coverage depth was ≥10. Mutations present in resistant but not parental sensitive sample (OVCAR4) were used to construct a phylogenetic tree accordingly using maximum parsimony. Nonsynonymous and synonymous exonic mutations were annotated using Annovar. Deep WGS data can be accessed at GitHub: https://github.com/WeiniHuangBW/OvarianCancerCellLines.

### Shallow whole genome sequencing (sWGS) and copy number analysis

100ng DNA was used for sWGS Libraries preparation with the ThruPLEX DNA-seq kit. Quality and quantity of the libraries were assessed with the DNA-7500 kit on a 2100 Bioanalyzer (Agilent Technologies) using a Kapa Library Quantification kit (Kapa Biosystems). Barcoded libraries were pooled together in equimolar amounts and each pool was sequenced on HiSeq4000 in SE-50bp mode.

Raw sequencing reads were checked for quality control (QC) using the FastQC program [25] and data found to be of good sequencing quality based on the phred scores. Reads were aligned against the reference human genome GRCh37 (hg19) using the bwa-mem method [26]. Post-alignment quality metrics were accessed using the qualimap program [27]. Duplicate reads were marked using the picard tools [28]. All bam files were sorted and indexed using samtools [29]. The R package QDNAseq [30] was used to call normalised segmented log2 copy numbers which were accessed and converted to absolute copy numbers using the R package ACE [31]. CN signatures were identified from the script provided [14]. Segmented data was annotated using the R package BiomaRt [32]. All analyses and plots were generated in R version 3.6.1.

### Statistical Analysis

Statistical analysis was performed using GraphPad Prism v7.04. Statistical significance was calculated using a two-tailed unpaired *t-*test unless otherwise specified. \**P*<0.05; ** *P*<0.001; *** *P*<0.0001.

## Results

### Platinum-resistance can be induced in HGSC cell lines and is maintained after drug withdrawal

To interrogate how platinum-resistance influences genomic and phenotypic evolution in HGSC, we first derived cisplatin- and carboplatin-resistant HGSC cells from three cell lines, OVCAR4, Ovsaho and Cov318, that had previously been categorised as “likely high grade serous” based on genomic features [17]. Cells were cultured *in vitro* in increasing concentrations of either cisplatin or carboplatin until an increase in IC_50_ between two and ten-fold was achieved to reflect the resistance observed in human patients [19]. Resistant cell lines (cisplatin-resistant: Ov4Cis, OvsahoCis and CovCis and carboplatin-resistant: Ov4Carbo and OvsahoCarbo) were then either cultured in the same drug dose (Ov4Cis: 0.6µm, OvsahoCis: 0.7µM, CovCis: 0.7µM, Ov4Carbo 3.5µM, OvsahoCarbo 5µM) or without drug (denoted ND) for eight weeks. IC_50_ was then measured again to determine if resistance had been maintained (**Fig 1A-C**). It was not possible to achieve additional resistance to carboplatin in Cov318 cells (data not shown) but in all other cases, resistance was induced successfully. In the OVCAR4-derived cells (Ov4Cis and Ov4Carbo), resistance to cisplatin and carboplatin was stable following drug withdrawal (**Fig 1A**). OvsahoCis cells also maintained resistance when drug was withdrawn (**Fig 1B**) but in OvsahoCarbo (**Fig 1B**) and CovCis (**Fig 1C**), resistance diminished over time after removal of drug from the culture medium (*P*<0.05). Ov4Cis (cisplatin-induced resistance) and Ov4Carbo (carboplatin-induced resistance) were cross-resistant to cisplatin and carboplatin with no significant difference in IC_50_ to either drug (**Fig 1D**).

**Figure 1:**
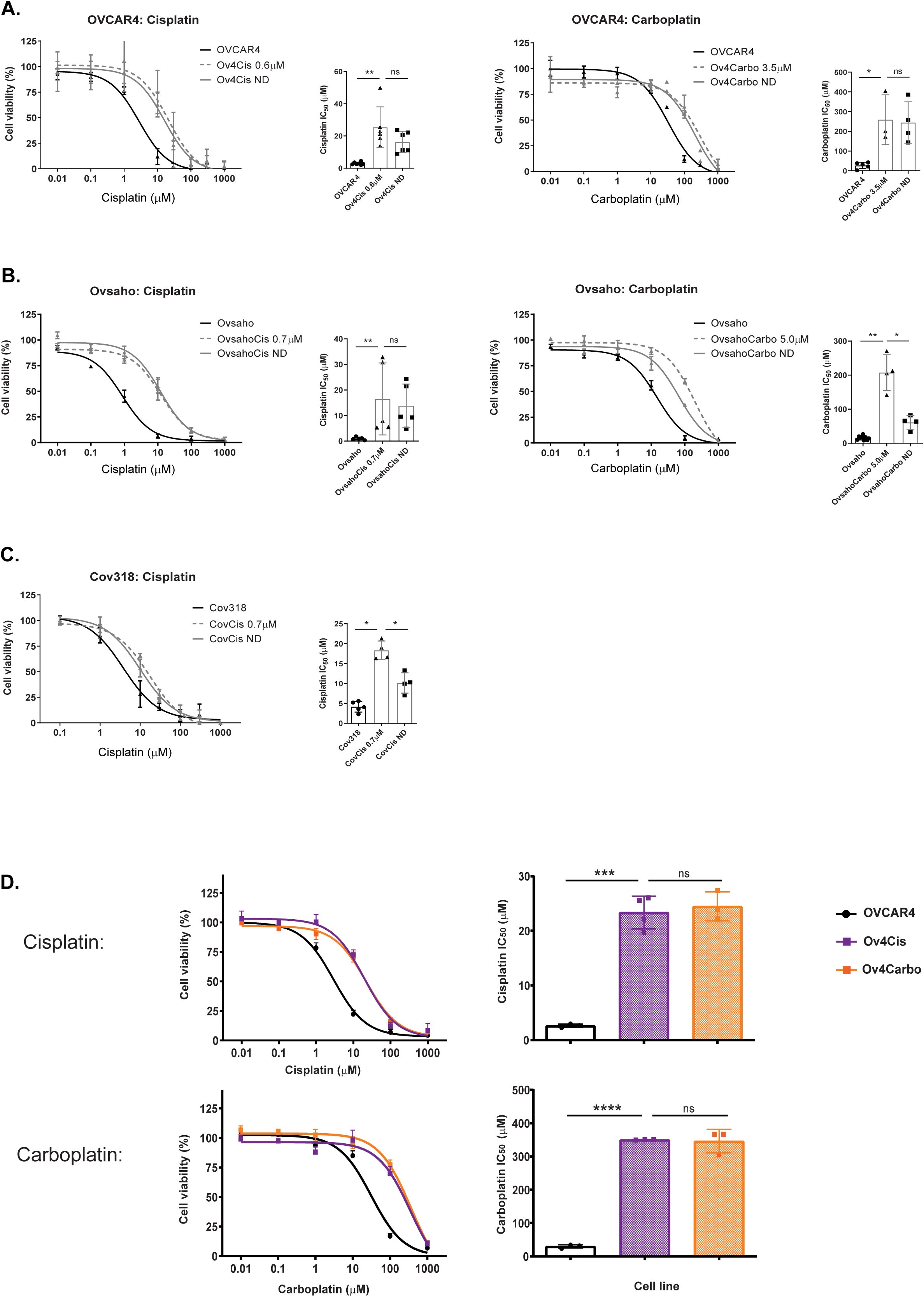
Platinum-resistance can be induced in HGSC cell lines and is maintained after drug withdrawal. OVCAR4 (**A**), Ovsaho (**B**) and Cov318 (**C**) cells were cultured in escalating doses of cisplatin or carboplatin. The resulting resistant cell lines (cisplatin-resistant: Ov4Cis, OvsahoCis and CovCis and carboplatin-resistant: Ov4Carbo and OvsahoCarbo) were then either cultured in the same drug dose (Ov4Cis: 0.6µm, OvsahoCis: 0.7µM, CovCis: 0.7µM, Ov4Carbo 3.5µM, OvsahoCarbo 5µM) or without drug (denoted ND) for eight weeks. Dose response and IC_50_ after 72 hours after drug treatment is shown. Black = sensitive cells, dashed grey = resistant cells, solid grey = resistant cells maintained without drug (ND). *N*=3-6, mean ± s.d., unpaired *t-*test, ns=non-significant, \**P*<0.05, \*\**P*<0.01. **D.** Dose response curves and IC_50_ to cisplatin and carboplatin 72 hours after drug administration. Black = OVCAR4, purple = Ov4Cis, orange = Ov4Carbo. *N*=3, mean ± s.d., unpaired *t-*test, ns = non-significant, \*\*\**P*<0.001, \*\*\*\**P*<0.0001.

### Platinum-resistant OVCAR4-derived clones, share multiple transcriptional features with platinum resistant/refractory human HGSC

Ov4Cis and Ov4Carbo cells were prioritised for further investigation because they maintained resistance when cultured without cisplatin and carboplatin respectively (**Fig 1A**). In addition, all three OVCAR4-derived cell lines (OVCAR4, Ov4Cis and Ov4Carbo) readily formed tumour spheroids when grown in non-adherent culture (**Suppl. Fig 1**) suggesting that they might form tumour nodules *in vivo*. To assess the similarity of these resistant cell lines to resistant human HGSC, we performed RNASeq on OVCAR4, Ov4Carbo and Ov4Cis. A total of 16,604 genes were sufficiently detected in our cell lines. Unsupervised clustering by principal component and hierarchical cluster analyses resulted in a clear segregation of the three sample groups indicating distinct transcriptomic features (**Suppl. Fig 2**). Differential expression analysis identified 1,763 genes in Ov4Carbo and 2,334 genes in Ov4Cis that had at least a two-fold change compared to OVCAR4 (**Suppl. Table 1**).

Next, gene expression differences between sensitive OVCAR4 cells and their resistant counterparts, Ov4Cis and Ov4Carbo, were compared to publicly available gene expression differences between whole tumour tissue obtained from patients with platinum-sensitive HGSC (n=31) versus patients with platinum-resistant and refractory HGSC (n=49) [24]. We observed approximately 500 genes in Ov4Cis (**Fig 2A**) and 330 genes in Ov4Carbo (**Fig 2B**) that had a concordant change with human samples (**Suppl. Table 2A**), demonstrating that transcriptional alterations induced by platinum-resistance in our models are also found in platinum-resistant human HGSC. Additional transcriptional changes were observed in human tumours, likely reflecting the microenvironmental response. We also identified changes in our models that were not detected or perhaps were missed in bulk tumour sequencing of human samples (**Figs 2A** and **B**). Pathway analysis of the concordant gene expression changes seen in both our resistant cell lines and human samples revealed that inflammatory and apoptotic pathways were overrepresented in down-regulated genes (**Fig 2C**) while extracellular matrix processes were overrepresented in up-regulated genes (**Fig 2D**) (Hypergeometric test, p<0.05, **Suppl. Table 2B-E**). Since these pathway changes occurred in Ov4Cis, Ov4Carbo and human samples they could represent unifying features that are relevant to multiple resistant settings.

**Figure 2:**
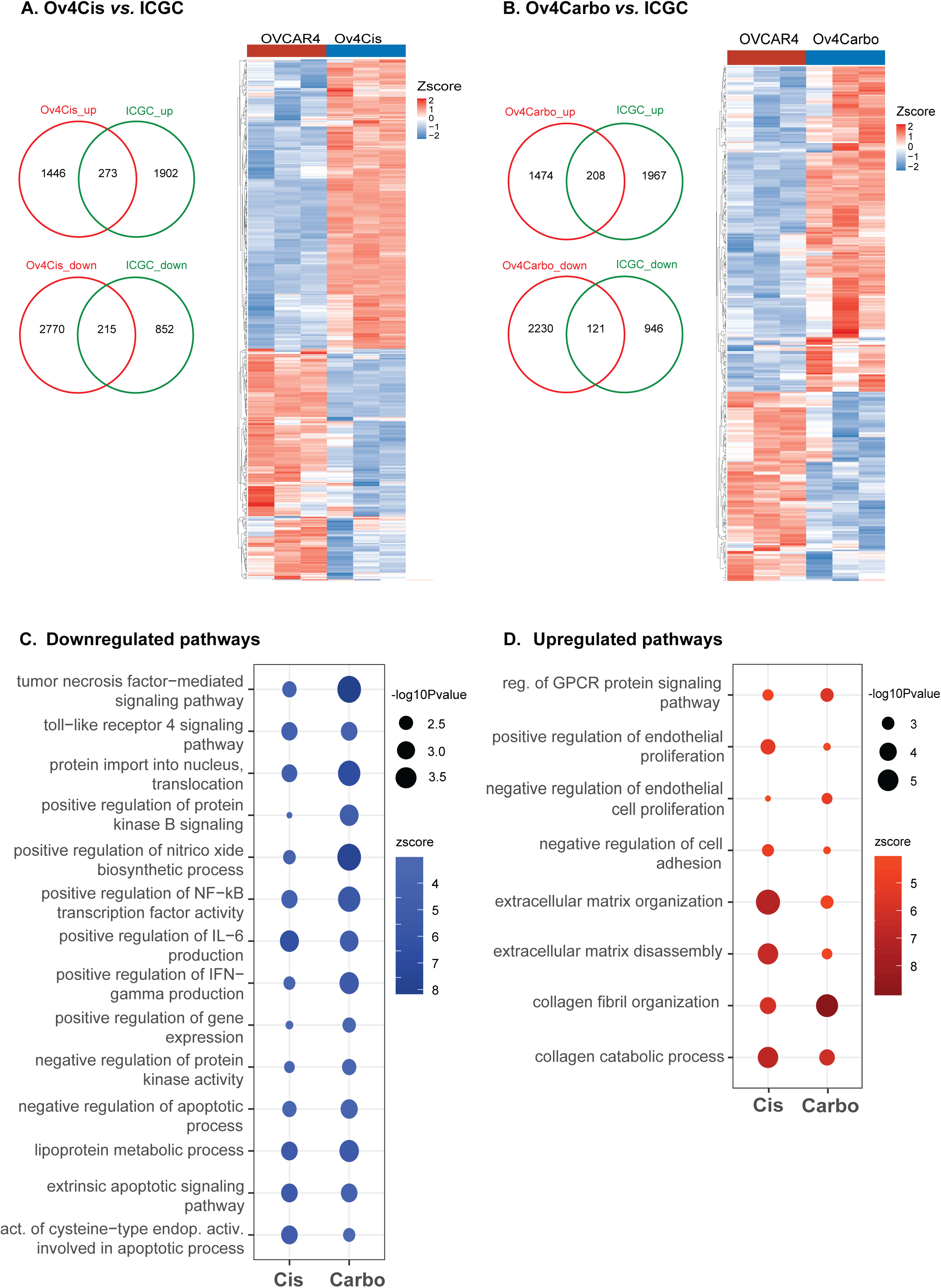
Platinum-resistant OVCAR4-derived clones, share multiple transcriptional features with platinum resistant/refractory human HGSC. Venn diagrams and heatmaps illustrating the differentially expressed protein-coding genes in **A.** Ov4Cis *vs.* OVCAR4 Parental and **B.** Ov4Carbo *vs*. OVCAR4 Parental and their overlap with gene expression differences between human resistant/refractory HGSC (N = 49) and sensitive HGSC (N = 31) obtained from the ICGC dataset. **C.** Concordantly down-regulated and **D.** concordantly up-regulated Gene Ontology Biological Processes (GOBP) in both resistant (Ov4Cis and Ov4Carbo) *vs.* sensitive (OVCAR4) cell lines and ICGC resistant/refractory *vs.* sensitive human HGSC are shown. Significance indicated by symbol size.

### Platinum resistance influences growth and invasion of subcutaneous tumours

To investigate the *in vivo* phenotype of these matched platinum-sensitive and resistant cells, tumours were established in both flanks of female nude mice (three mice per cell line) and subcutaneous (SC) tumours were measured weekly using callipers (**Fig 3A**). All cell lines were able to form SC tumours but tumour growth varied markedly between the resistant cells, so that the growth rate (cm^2^/week) of Ov4Cis was significantly lower than that of Ov4Carbo (*P*<0.05, **Fig 3B**). HGSC histology was confirmed in all tumours by H+E staining and by positive IHC for mutant p53 and the Mullerian transcription factor, PAX8 (**Fig 3C**). Interestingly, both H+E and IHC showed that Ov4Cis cells were densely packed within smaller SC tumours, whereas in OVCAR4 and Ov4Carbo tumours, malignant cells were more dispersed and stromal components more prominent. Together, these observations suggest that, despite their comparable resistance to cisplatin and carboplatin, Ov4Cis had evolved a less invasive phenotype in response to platinum treatment, while Ov4Carbo cells had acquired more aggressive *in vivo* characteristics.

**Figure 3:**
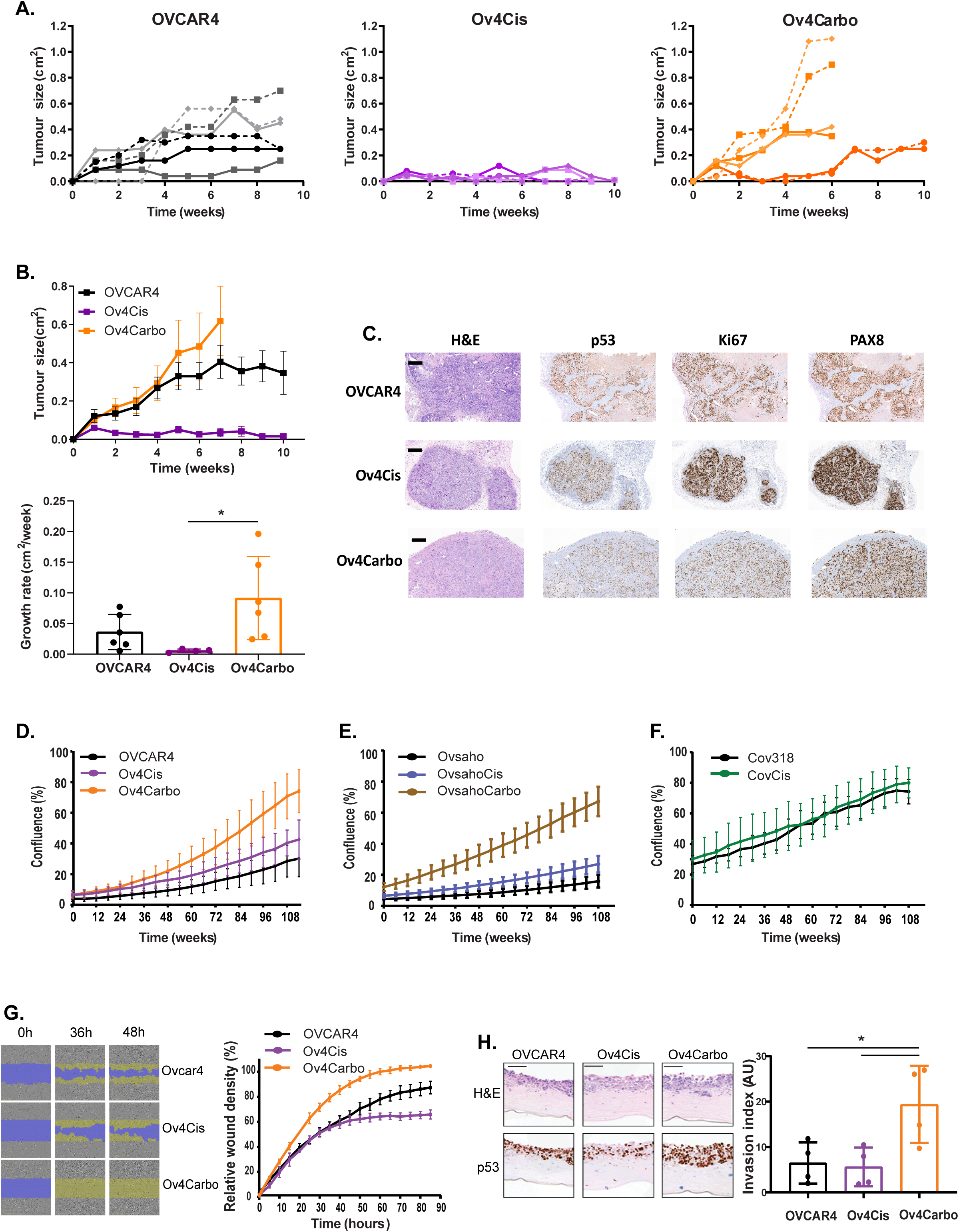
The evolution of platinum resistance influences growth and invasion of subcutaneous tumours. **A.** SC xenografts of OVCAR4, Ov4Cis or Ov4Carbo in both flanks of female CD1*nu/nu* mice (*N*=3 mice per cell line). Black = OVCAR4, purple = Ov4Cis, orange = Ov4Carbo. **B.** Size by calliper measurement (cm^2^) and growth rate (cm^2^/week) of SC OVCAR4, Ov4Cis and Ov4Carbo xenografts. *N*=4-6 tumours per model, mean ± s.d., one-way ANOVA \**P*<0.05. **C.** Haemotoxylin and eosin staining (H+E) staining showing greater tumour cell density in Ov4Cis tumour nodules compared to OVCAR4 and Ov4Carbo. Immunohistochemistry (IHC) demonstrating positive staining for p53, the Mullerian transcription factor PAX8 and Ki67. Scale bars = 500µm. **D.** Proliferation of OVCAR4, Ov4Cis and Ov4Carbo, **E.** Ovsaho, OvsahoCis and OvsahoCarbo and **F**. Cov318 and CovCis over time. **G.** Migration of OVCAR4, Ov4Cis and Ov4Carbo cells measured over time after scratch formation. Representative images are shown at 0, 36 and 48 hours. **H.** Invasion of OVCAR4, Ov4Cis and Ov4Carbo cells co-cultured with human HGSC omental fibroblasts through Matrigel/collagen gels. Invasion index = cell number x area X maximum depth. Black = OVCAR4, purple = Ov4Cis, orange = Ov4Carbo. *N*=3, mean ± s.d., unpaired *t-*test, \**P*<0.05. Representative images at day 9 are shown. Scale bars = 200µm.

We then investigated *in vitro* behaviours that could explain these diverse *in vivo* phenotypes. Ov4Carbo cells were found to proliferate more rapidly than OVCAR4 *in vitro* (**Fig 3D**). Interestingly, the same pattern was observed in Ovsaho-derived cells where again, the carboplatin-resistant cell line, OvsahoCarbo proliferated more rapidly than parental Ovsaho cells (**Fig 3E**). In contrast, cisplatin resistance did not alter the proliferation of any cell line tested (Ov4Cis, OvsahoCis, CovCis, **Fig 3D-F**). Ov4Carbo cells also migrated more efficiently (**Fig 3G**) and when co-cultured with human HGSC omental fibroblasts in three dimensional organotypic cultures, Ov4Carbo cells were significantly more invasive than both Ov4Cis and OVCAR4 (*P*<0.05) (**Fig 3H**).

### Platinum-resistant HGSC evolves diverse intraperitoneal phenotypes

To further explore this phenotypic diversity in platinum-resistant HGSC, we used a more clinically-relevant orthotopic, intraperitoneal (IP) system. First, the OVCAR4-derived cell panel were transfected with lentiviruses expressing Firefly luciferase (Luc) to create OVCAR4-Luc, Ov4Cis-Luc and Ov4Carbo-Luc. *In vitro* resistance to cisplatin and carboplatin was maintained in these transfected cells (**Suppl. Fig 3**). IP xenografts were then created in female nude mice. Mice were monitored with weekly bioluminescence imaging (BLI) and killed when they reached humane endpoints pre-defined by the UK Home Office. Median survival of mice bearing OVCAR4-Luc was 100 days. In keeping with the increased growth rate of carboplatin-resistant Ov4Carbo as SC xenografts (**Fig 3B**), Ov4Carbo-Luc cells had a more aggressive IP phenotype with a reduced median survival of 56 days. In contrast, but also in-line with the reduced invasive ability of Ov4Cis cells as SC xenografts, no mice with IP Ov4Cis-Luc xenografts had reached Home Office endpoint at 140 days (log-rank test *P*<0.001) (**Fig 4A**). These survival differences were reflected in BLI in individual mice over time (**Fig 4B**).

**Figure 4:**
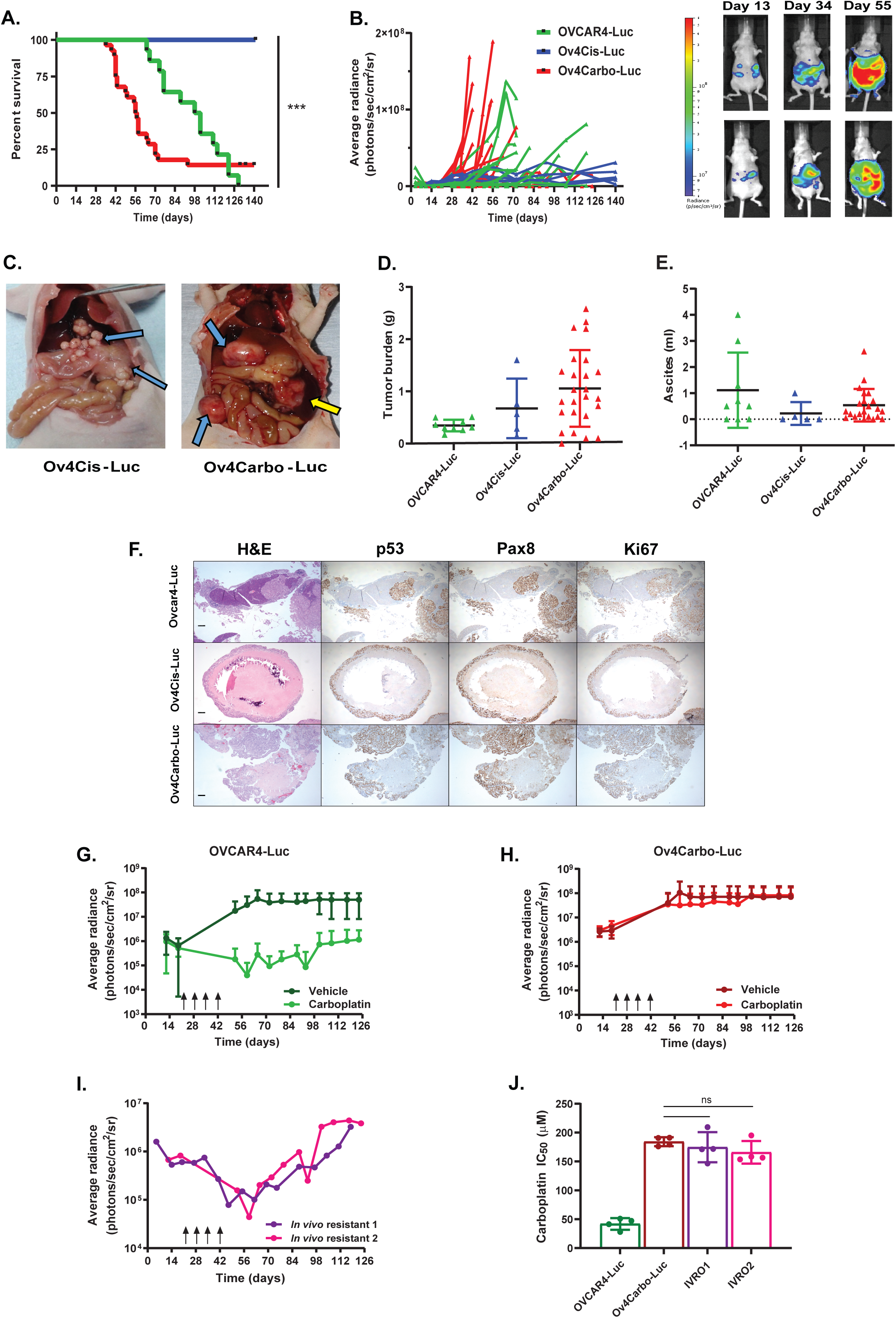
Platinum-resistant HGSC evolves diverse intraperitoneal phenotypes. **A.** IP OVCAR4-Luc, Ov4Cis-Luc or Ov4Carbo-Luc xenografts were created in female CD1*nu/nu* mice. Green = OVCAR4-Luc (*N*=14), red = Ov4Carbo-Luc (*N*=28), blue = Ov4Cis-Luc (*N*=5) \*\*\**P*<0.001, log-rank test. **B.** Corresponding light output (BLI) and representative images (Ov4Carbo-Luc) are shown over time. **C.** Macroscopic images in Ov4Cis-Luc and Ov4Carbo-Luc xenograft taken at necropsy. Ov4Cis-Luc xenografts remained localised, creating multiple small nodules (blue arrows) without invasion of other organs. Ov4Carbo-Luc produced multiple peritoneal nodules (blue arrows) and haemorrhagic ascites (yellow arrows). **D.** Tumour weight (g) and **E.** ascites volume (ml) at necropsy. Mean ± s.d. **F.** Pathological examination (H+E and IHC) demonstrating positive staining for p53, the Mullerian transcription factor PAX8, and Ki67 in all three models. Scale bars = 200µm. **G and H.** Female CD1*nu/nu* mice with intraperitoneal (IP) xenografts of either OVCAR4-Luc or Ov4Carbo-luc cells. Mice were treated IP with either carboplatin 50mg/kg or vehicle control on days 21, 28, 35 and 42 (black arrows). Average radiance (photons/sec/cm^2^/sr) is shown over time. **G.** Dark green = vehicle-treated OVCAR4-Luc xenografts, light green = carboplatin-treated OVCAR4-Luc xenografts, **H.** dark red = vehicle-treated Ov4Carbo-Luc xenografts, light red = carboplatin-treated Ov4Carbo-Luc xenografts. *N*=4-5 per group, mean ± s.d. **I.** At Home Office end point OVCAR4-Luc tumours were harvested from two separate mice that had both received carboplatin. Light output from these two mice over time is shown. Tumours were homogenized and cells cultured *in vitro* to create the carboplatin-resistant cell lines, IVR01 (purple: *in vivo* resistant 1) and IVR02 (pink: *in vivo* resistant 2). **J.** IC_50_ to carboplatin at 72 hours. *N*=4, mean ± s.d., ns = non-significant.

Post-mortem examinations were conducted on all mice at end point and revealed that OVCAR4-Luc (not shown) and Ov4Carbo-Luc IP tumours (**Fig 4C**) mirrored the human disease with multiple peritoneal nodules and ascites. In keeping with their reduced survival, Ov4Carbo-Luc xenografts had the highest tumour burden at necropsy, although there was wide variation in measured tumour weight between individual mice (**Fig 4D**). Interestingly, although tumour weight in Ov4Cis-Luc xenografts was comparable to the other models (**Fig 4D**), these Ov4Cis-Luc tumour nodules remained discrete without invasion of nearby structures (**Fig 4C**) or formation of ascites (**Fig 4E**). Additionally, pathological examination revealed that Ov4Cis-Luc nodules consisted of a rim of tumour tissue with a central necrotic core while Ov4Carbo-Luc tumours were locally invasive (**Fig 4F**). These discoveries further endorse our previous observation of diverse evolutionary trajectories during resistance to platinum therapy, with Ov4Cis cells developing a markedly less invasive phenotype, while Ov4Carbo cells evolve more aggressive clinical behaviour.

We next explored the impact of inducing platinum resistance *in vivo*. Again female nude mice were injected IP with either OVCAR4-Luc (**Fig 4G**) or Ov4Carbo-Luc cells (**Fig 4H**). Once tumours had established at day 22, mice were treated IP with weekly injections of either 50mg/kg carboplatin or vehicle alone for four weeks. Tumour growth was monitored with BLI. As expected, Ov4Carbo-Luc cells were resistant to carboplatin treatment and light emission overlapped with vehicle-treated control mice (**Fig 4H**). In contrast, mice with OVCAR4-Luc IP xenografts initially responded to carboplatin but tumours eventually regrew indicating the *in vivo* development of carboplatin-resistance (**Fig 4G**). The IP OVCAR4-Luc tumours of two carboplatin-treated mice were harvested at end point, (**Fig 4I**) and the resulting cell lines were denoted IVR01 and IVR02 (*in vivo* resistant 1 and 2). *In vitro* (Ov4Carbo) and *in vivo* (IVR01 and IVR02) -derived carboplatin-resistant cells had equivalent IC_50_ to carboplatin (all *P*<0.0001 compared to OVCAR4) (**Fig 4J**), despite the different strategies used to induce resistance. Since IVR01 and IVR02 were harvested 12 weeks after the last platinum treatment, we considered that resistance was stable after drug withdrawal in these cell lines.

### Divergent *in vivo* phenotypes are explained by copy number and gene expression changes rather than specific mutations

To interrogate these distinct *in vivo* phenotypes we conducted deep whole genome sequencing on OVCAR4, Ov4Cis and four separate clones grown from single Ov4Cis cells (clones 1, 2, 3 and 4). Variant point mutations were identified by comparing to the human reference genome (hg19). Mutations present in resistant samples but not OVCAR4 cells were then selected and used to construct a phylogenetic tree (**Fig 5A**). Only two nonsynonymous exonic mutations were shared between all five resistant cells; FCGBP (Fc Fragment Of IgG Binding Protein) and RTL1 (Retrotransposon Gag Like 1). A total of 161 nonsynonymous mutations separated Ov4Cis from OVCAR4. By overlaying these 161 mutant genes with our previous RNASeq analysis, we found that 38 of these 161 genes were also differentially expressed between OVCAR4 and Ov4Cis (**Suppl. Table 3**).

**Figure 5:**
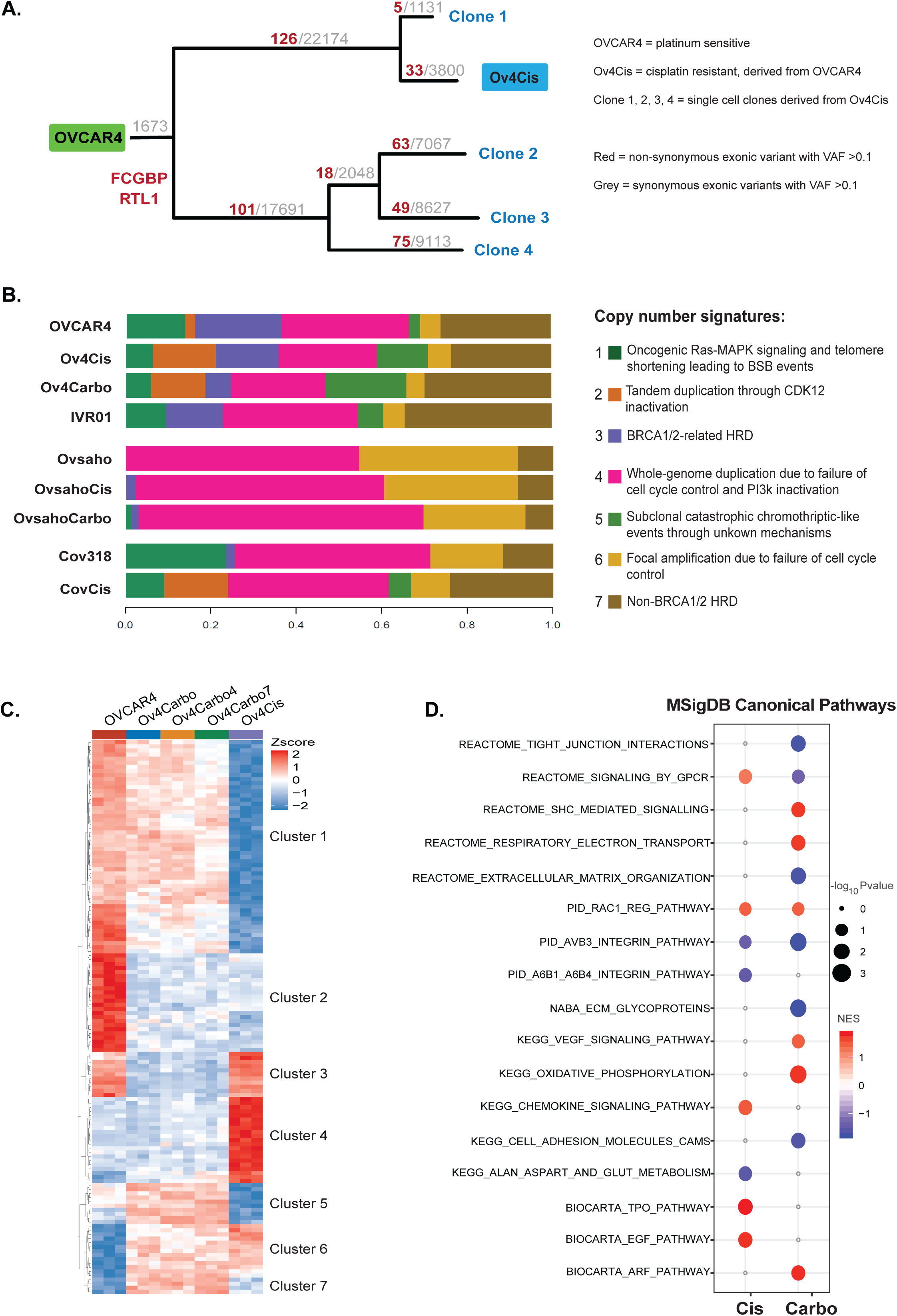
Divergent *in vivo* phenotypes are explained by copy number and gene expression changes rather than specific mutations. **A.** Deep whole genome sequencing (50x mean depth, Illumina’s HiSeq X Ten) of OVCAR4, Ov4Cis and four cell lines derived from single Ov4Cis clones (clones 1-4). Red = number of non-synonymous exonic variant with variant allele frequency >0.01, grey = number of synonymous exonic variant with variant allele frequency >0.01. **B.** Shallow whole genome sequencing of all cell lines as listed in figure. Copy number signatures were called as we previously described. Seven copy number signatures are colour coded as indicated and relative proportions for each cell line are shown **C.** RNASeq (32x mean depth, Illumina HiSeq4000) of OVCAR4, Ov4Cis and Ov4Carbo as well as two cell lines derived from single cell Ov4Carbo clones (Ov4Carbo4 and Ov4Carbo7). Heatmap illustrating expression of genes that maximally separate the three major classes of cell lines; sensitive (OVCAR4), carboplatin resistant (Ov4Carbo, Ov4Carbo4 and Ov4Carbo7) and cisplatin resistant (Ov4Cis) (FDR = 0.001). **D.** Significantly up-regulated (red) or down-regulated (blue) GSEA canonical pathways in platinum-resistant Ov4Cis and Ov4Carbo cells compared to sensitive OVCAR4 cells. Significance indicated by symbol size.

We then applied our newly defined copy number signatures [14] to our entire cell panel (**Fig 5B**). Ancestral, sensitive cell lines each displayed a unique profile, which changed during the evolution of platinum resistance. Although these changes appeared to be specific to each resistant cell line, recurrent, resistance-associated changes were observed so that in comparison to matched sensitive cells, signatures 1, 3 and 6 tended to decrease in resistant cells, while signatures 2, 5 and 7 tended to increase. To interrogate the potential functional implications of these changes, we compared the copy number alterations (CNAs) in the two resistant OVCAR4-derived cell lines (Ov4Cis and Ov4Carbo) with parental sensitive OVCAR4 cells and again overlaid these resistance-associated CNAs with our RNASeq analysis. Between 8 and 18% of CNAs were reflected in the gene expression comparisons (**Suppl. Fig 4** and individual genes shown in **Suppl. Table 4**). Further analysis of these overlapping features revealed that DNA Recombination Pathway was increased in both OV4Cis (*P*=0.0036) and Ov4Carbo (*P*=0.00026) resistant cell lines (**Suppl Table 5**, yellow highlight) and that expression of Rad51C (OV4Cis and Ov4Carbo), Rad51D (OV4Cis) and BRCA1 (OV4Cis and Ov4Carbo) were all increased compared to OVCAR4 cells (**Suppl. Table 4** yellow highlight). These changes could potentially account for the reduction in signature 3 (BRCA1/2-related HRD and good overall survival) we observed in these chemotherapy-resistant cells. Moreover, Rad51D expression was increased in the ICGC human resistant/refractory dataset, implying that similar changes also occur in human patients.

Gene expression differences (RNASeq) associated with platinum resistance were then interrogated in greater depth in OVCAR4, Ov4Cis and Ov4Carbo cells as well as two lines that we had derived from single cell Ov4Carbo clones (Ov4Carbo4 and Ov4Carbo7) (**Fig 5C**). Analysis of genes that maximally separated the different cell lines (FDR 0.001) revealed seven distinct gene clusters (most significant are shown in **Fig 5C** and **Suppl. Table 6**, full list **Suppl. Table 7**). Clusters 2 and 6 separated OVCAR4 from all resistant cells: Ov4Cis, Ov4Carbo, Ov4Carbo4 and Ov4Carbo7. Rather than directing phenotype, these gene expression changes are likely to be responsible for shared mechanisms of drug resistance and the cross-resistance to cisplatin and carboplatin that we previously observed (**Fig 1D** and **Suppl. Fig 3**). In keeping with their transcriptomic similarity to human HGSC (**Fig 2**), ABCB1, a known multi-drug resistance gene that is frequently induced by chemotherapy resistance in human HGSC [33], was also strongly up-regulated in our resistant cell lines (LogFc 8.78, **Suppl Table 7**, yellow highlight).

In contrast, other distinct clusters, (clusters 1, 3, 4, 5, 7) separated the Ov4Carbo-derived cell lines from Ov4Cis and thus likely account for the diverse *in vitro* and *in vivo* behaviour we observed. Gene set enrichment analysis (GSEA, **Fig 5D** and **Suppl. Table 8**) provided more detail on these features. Only two pathways were concordantly regulated in resistant cells compared to OVCAR4 (PID_ RAC_ REG_ PATHWAY and PID_AVB3_INTEGRIN_PATHWAY). In contrast, several pathways were identified that could explain the diverse behaviours seen in resistant cells. For example, pathways associated with oncogenesis, invasion and metastasis including REACTOME_ SHC_ MEDIATED_ SIGNALLING, REACTOME_ RESPIRATORY_ ELECTRON_ TRANSPORT, KEGG_ VEGF_ SIGNALING_ PATHWAY and KEGG_ OXIDATIVE_ PHOSPHORYLATION, were selectively increased in Ov4Carbo compared to OVCAR4 cells, while pathways involved in cell adhesion and integrity of the extracellular matrix such as REACTOME_ TIGHT_ JUNCTION_ INTERACTIONS, REACTOME_ EXTRACELLULAR_ MATRIX_ ORGANIZATION, NABA_ ECM_ GLYCOPROTEINS and KEGG_ CELL_ ADHESION_ MOLECULES_ CAMS were down-regulated in Ov4Carbo cells. Together, these marked differences in gene expression between resistant cell lines likely account for the more migratory and locally invasive behaviour of Ov4Carbo cells. These clear transcriptional changes, rather than genetic mutations, therefore appear to mediate the diverse platinum-resistant phenotypes we have observed and provide a new functional context for the copy number signatures we previously defined in human HGSC.

## Discussion

Drug resistance remains a major obstacle in cancer care. Research to address this has focused predominantly on the genetic and biochemical changes associated with resistance to specific anti-cancer agents [13, 34, 35]. One of the most comprehensive investigations into chemotherapy-resistant HGSC revealed a variety of genetic mediators of platinum-resistance including increased expression of the ABCB1 drug efflux pump, a change that was reproduced in our resistant cell lines, as well as reversion mutations in BRCA genes and loss of BRCA promoter-methylation [13]. Unfortunately this wealth of molecular information has not yet lead to effective new therapies. The recent appreciation that cancer is subject to clonal selection and Darwinian evolution [36] provides a new lens with which to view the pervasive clinical problem of drug resistance. In this evolutionary model, cancer cells with a growth advantage in the presence of therapy will expand to create drug-resistant tumours [37]. This means that selection ultimately acts on phenotypic variation [38]. Diverse clinical phenotypes and disease trajectories are observed in patients with relapsed HGSC [39] but to date, there has been little characterisation of the biological changes that underpin these behaviours.

Our novel platinum-resistant HGSC models provide new insights into the evolution of resistance and are likely to be clinically informative because they share genetic and transcriptional profiles with platinum-resistant human HGSC. Moreover, they accurately reproduce the phenotypic diversity seen in patients. The infiltrative and metastatic intraperitoneal phenotype produced by Ov4Carbo cells is analogous to the most usual pattern of recurrent, human HGSC [40]. However, an oligometastatic picture, focused on localised disease, lymph node metastasis and a favourable outcome also occurs clinically [41] and Ov4Cis cells are a useful model for this more benign disease course. Despite the phenotypic diversity that we observed, resistance to cisplatin and carboplatin was comparable in the different cell lines, regardless of the drug initially used to induce resistance and whether resistance was induced *in vitro* or *in vivo.* This shared resistance can be explained by our RNASeq and copy number analysis, both of which identified common changes in multiple different resistant clones, demonstrating that a small number of context-independent alterations were responsible for therapy resistance.

The frequency of recurrent oncogenic mutations in HGSC is low and in keeping with this, our WGS identified few mutational changes associated with cisplatin resistance. Only two nonsynonymous mutations were common to all five of the cisplatin-resistant cells tested and to our knowledge, neither have been previously linked to drug resistance. In other cancer types, multiple genetic mechanisms of resistance have been shown to evolve simultaneously [42, 43] despite the fact that a cell may only require a small number of these to establish resistance [44]. This diversity appears to have an evolutionary advantage and in breast cancer, a greater variety of transcriptional resistance mechanisms results in enhanced metastatic potential [45]. We also observed multiple gene expression changes in our resistant cells and these complex, transcriptional changes that are unique to individual, resistant cells, are likely to be responsible for the phenotypic diversity we observed.

Early and ubiquitous mutation of TP53 likely drives the chromosomal instability that characterises HGSOC [14]. This genetic complexity provides cells with abundant adaptive potential but has been challenging to model and characterise. To address this, we previously categorised HGSC genomes into seven CN signatures [14] but how these change during platinum resistance has not previously been investigated. By applying the same approach to our cell panel, we show here that, like human HGSC, our cell lines each display a mixture of signatures with different relative frequencies in the three ancestral sensitive HGSC cell lines. Importantly, this study provides the first evidence that CN profiles can change during the evolution of platinum resistance. Although these changes appeared to be cell-line specific, certain patterns were consistent across our cell panel. Signatures 1, 3 and 6 decreased whereas signatures 2, 5, and 7 increased and it is possible that this combination of signatures provides a fitness advantage during platinum therapy. We previously demonstrated that CN signatures correlate with patient prognosis [14]. By overlapping CN changes with RNASeq data, we were able to demonstrate specific and relevant changes in gene expression (e.g. increased Rad51C) that could potentially mediate these clinical behaviours. Further functional interrogation of these CN changes and correlation with human samples, may provide new insights into platinum-resistant HGSC.

In summary, we have defined and characterised a unique panel of clinically relevant and usable, platinum-resistant HGSC models that are now freely available. We have discovered that specific transcriptional alterations can induce resistance in multiple disease contexts. These co-exist with a distinct repertoire of transcriptional changes that mediate phenotypic diversity, echoing human cancer. We have carried out the first mechanistic investigation into our newly characterised copy number signatures and reveal for the first time that these unique profiles undergo specific, recurrent alterations during drug therapy. Since we have shown that these resistance-associated CNAs correlate with relevant changes in gene expression, we anticipate they could potentially reveal new therapeutic opportunities in platinum-resistant HGSC.

## Acknowledgements

We thank Ashley Browne for originally creating OVCAR4-Luc, Ov4Cis, Ov4Cis-Luc, Ov4Carbo, Ov4Carbo-Luc, CovCis and OvsahoCis, Jacqueline McDermott for interpretation of pathology slides, Cheng Zhao and Ann-Marie Baker for experimental and logistic contributions to shallow and deep whole genome sequencing respectively and to Prof Tyson Sharp and Dr Sarah Martin for reviewing and advising on this manuscript.

## References

1. McCluggage, W.G., Morphological subtypes of ovarian carcinoma: a review with emphasis on new developments and pathogenesis. Pathology, 2011. 43(5): p. 420–32.

2. Bowtell, D.D., et al., Rethinking ovarian cancer II: reducing mortality from high-grade serous ovarian cancer. Nat Rev Cancer, 2015. 15(11): p. 668–79.

3. UK, C.R. https://www.cancerresearchuk.org/health-professional/cancer-statistics/statistics-by-cancer-type/ovarian-cancer/survival#heading-Three. [cited 2019.

4. Ahmed, A.A., et al., Driver mutations in TP53 are ubiquitous in high grade serous carcinoma of the ovary. J Pathol, 2010. 221(1): p. 49–56.

5. Tothill, R.W., et al., Novel molecular subtypes of serous and endometrioid ovarian cancer linked to clinical outcome. Clin Cancer Res, 2008. 14(16): p. 5198–208.

6. Cancer Genome Atlas Research, N., Integrated genomic analyses of ovarian carcinoma. Nature, 2011. 474(7353): p. 609–15.

7. Ledermann, J., et al., Olaparib maintenance therapy in platinum-sensitive relapsed ovarian cancer. N Engl J Med, 2012. 366(15): p. 1382–92.

8. Coleman, R.L., et al., Rucaparib maintenance treatment for recurrent ovarian carcinoma after response to platinum therapy (ARIEL3): a randomised, double-blind, placebo-controlled, phase 3 trial. Lancet, 2017. 390(10106): p. 1949–1961.

9. Mirza, M.R., et al., Niraparib Maintenance Therapy in Platinum-Sensitive, Recurrent Ovarian Cancer. N Engl J Med, 2016. 375(22): p. 2154–2164.

10. Moore, K., et al., Maintenance Olaparib in Patients with Newly Diagnosed Advanced Ovarian Cancer. N Engl J Med, 2018. 379(26): p. 2495–2505.

11. Gonzalez-Martin, A., et al., Niraparib in Patients with Newly Diagnosed Advanced Ovarian Cancer. N Engl J Med, 2019.

12. Coleman, R.L., et al., Veliparib with First-Line Chemotherapy and as Maintenance Therapy in Ovarian Cancer. N Engl J Med, 2019.

13. Patch, A.M., et al., Whole-genome characterization of chemoresistant ovarian cancer. Nature, 2015. 521(7553): p. 489–94.

14. Macintyre, G., et al., Copy number signatures and mutational processes in ovarian carcinoma. Nat Genet, 2018. 50(9): p. 1262–1270.

15. Ince, T.A., et al., Characterization of twenty-five ovarian tumour cell lines that phenocopy primary tumours. Nat Commun, 2015. 6: p. 7419.

16. Elias, K.M., et al., Beyond genomics: critical evaluation of cell line utility for ovarian cancer research. Gynecol Oncol, 2015. 139(1): p. 97–103.

17. Domcke, S., et al., Evaluating cell lines as tumour models by comparison of genomic profiles. Nat Commun, 2013. 4: p. 2126.

18. Mitra, A.K., et al., In vivo tumor growth of high-grade serous ovarian cancer cell lines. Gynecol Oncol, 2015. 138(2): p. 372–7.

19. McDermott, M., et al., In vitro Development of Chemotherapy and Targeted Therapy Drug-Resistant Cancer Cell Lines: A Practical Guide with Case Studies. Front Oncol, 2014. 4: p. 40.

20. Jenei, V., M.L. Nystrom, and G.J. Thomas, Measuring invasion in an organotypic model. Methods Mol Biol, 2011. 769: p. 223–32.

21. Hansen, K.D., R.A. Irizarry, and Z. Wu, Removing technical variability in RNA-seq data using conditional quantile normalization. Biostatistics, 2012. 13(2): p. 204–16.

22. Ritchie, M.E., et al., limma powers differential expression analyses for RNA-sequencing and microarray studies. Nucleic Acids Res, 2015. 43(7): p. e47.

23. Subramanian, A., et al., Gene set enrichment analysis: a knowledge-based approach for interpreting genome-wide expression profiles. Proc Natl Acad Sci U S A, 2005. 102(43): p. 15545–50.

24. ICGC. ICGC Data Portal. 2019 [cited 2019; Available from: http://dcc.icgc.org.

25. Andrews, S. FastQC: a quality control tool for high throughput sequence data. 2010 [cited 2020; Available from: http://www.bioinformatics.babraham.ac.uk/projects/fastqc.

26. H., L., Aligning sequence reads, clone sequences and assembly contigs with BWA-MEM. 1303.3997v1 [q-bio.GN], 2013.

27. Okonechnikov, K., A. Conesa, and F. Garcia-Alcalde, Qualimap 2: advanced multi-sample quality control for high-throughput sequencing data. Bioinformatics, 2016. 32(2): p. 292–4.

28. Broad Institute, G.R. Picard Toolkit. 2019; Available from: http://broadinstitute.github.io/picard/.

29. Li, H., et al., The Sequence Alignment/Map format and SAMtools. Bioinformatics, 2009. 25(16): p. 2078–9.

30. Scheinin, I., et al., DNA copy number analysis of fresh and formalin-fixed specimens by shallow whole-genome sequencing with identification and exclusion of problematic regions in the genome assembly. Genome Res, 2014. 24(12): p. 2022–32.

31. Poell, J.B., et al., ACE: absolute copy number estimation from low-coverage whole-genome sequencing data. Bioinformatics, 2019. 35(16): p. 2847–2849.

32. Durinck, S., et al., Mapping identifiers for the integration of genomic datasets with the R/Bioconductor package biomaRt. Nat Protoc, 2009. 4(8): p. 1184–91.

33. Christie, E.L., et al., Multiple ABCB1 transcriptional fusions in drug resistant high-grade serous ovarian and breast cancer. Nat Commun, 2019. 10(1): p. 1295.

34. Siddik, Z.H., Cisplatin: mode of cytotoxic action and molecular basis of resistance. Oncogene, 2003. 22(47): p. 7265–79.

35. Nikolaou, M., et al., The challenge of drug resistance in cancer treatment: a current overview. Clin Exp Metastasis, 2018. 35(4): p. 309–318.

36. Greaves, M. and C.C. Maley, Clonal evolution in cancer. Nature, 2012. 481(7381): p. 306–13.

37. Gatenby, R.A., J. Brown, and T. Vincent, Lessons from applied ecology: cancer control using an evolutionary double bind. Cancer Res, 2009. 69(19): p. 7499–502.

38. Gillies, R.J., D. Verduzco, and R.A. Gatenby, Evolutionary dynamics of carcinogenesis and why targeted therapy does not work. Nat Rev Cancer, 2012. 12(7): p. 487–93.

39. Pignata, S., et al., Treatment of recurrent ovarian cancer. Ann Oncol, 2017. 28(suppl_8): p. viii51–viii56.

40. Amate, P., et al., Ovarian cancer: sites of recurrence. Int J Gynecol Cancer, 2013. 23(9): p. 1590–6.

41. Hollis, R.L., et al., Clinical and molecular characterization of ovarian carcinoma displaying isolated lymph node relapse. Am J Obstet Gynecol, 2019. 221(3): p. 245 e1–245 e15.

42. Cree, I.A. and P. Charlton, Molecular chess? Hallmarks of anti-cancer drug resistance. BMC Cancer, 2017. 17(1): p. 10.

43. Shi, H., et al., Acquired resistance and clonal evolution in melanoma during BRAF inhibitor therapy. Cancer Discov, 2014. 4(1): p. 80–93.

44. Tegze, B., et al., Parallel evolution under chemotherapy pressure in 29 breast cancer cell lines results in dissimilar mechanisms of resistance. PLoS One, 2012. 7(2): p. e30804.

45. Nguyen, A., et al., Highly variable cancer subpopulations that exhibit enhanced transcriptome variability and metastatic fitness. Nat Commun, 2016. 7: p. 11246.

