## supplemental figures and legends for "Platinum resistance induces diverse evolutionary trajectories in high grade serous ovarian cancer"

#### Supplemental Figure 1

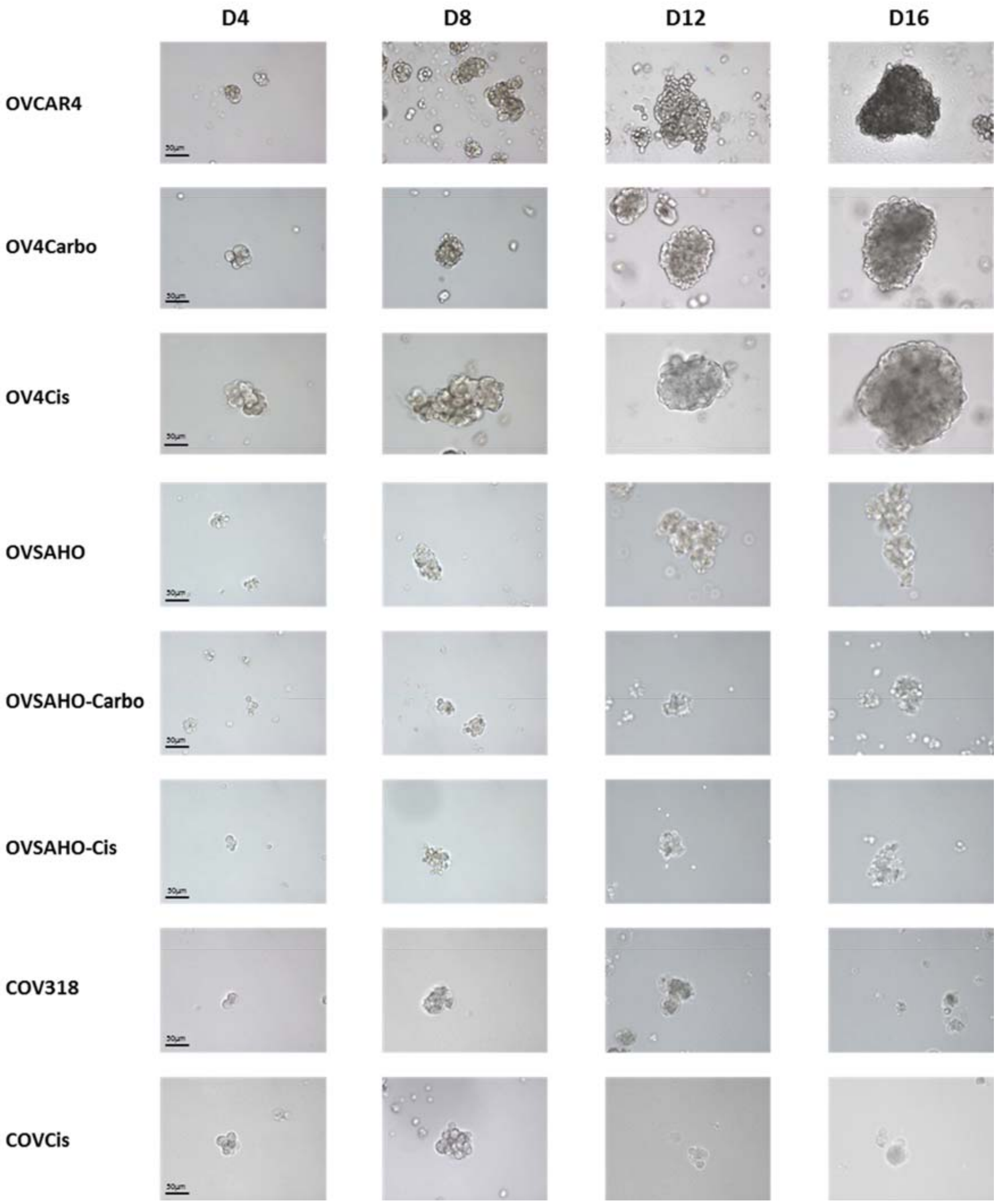

### Supplemental Figure 2

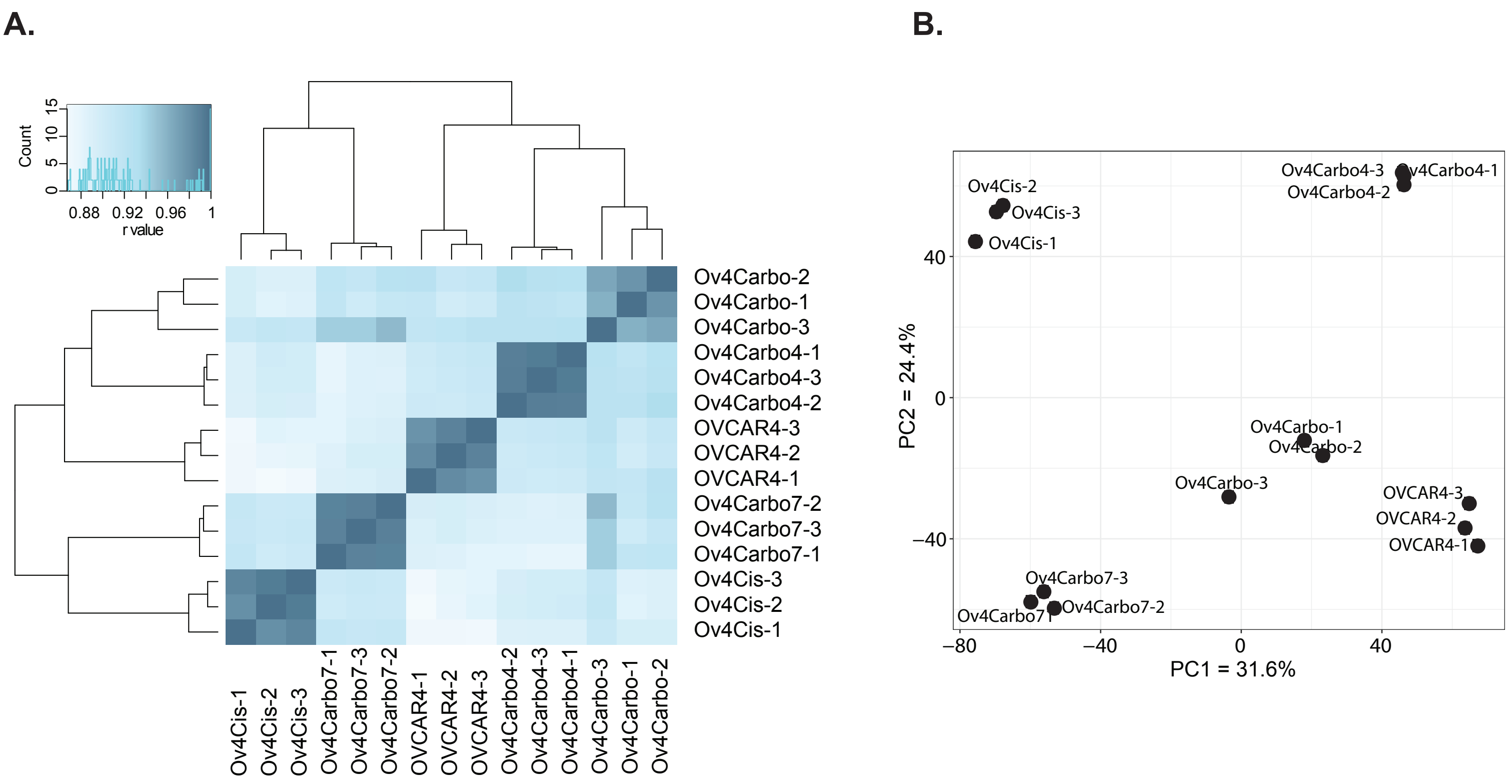

### Supplemental Figure 3

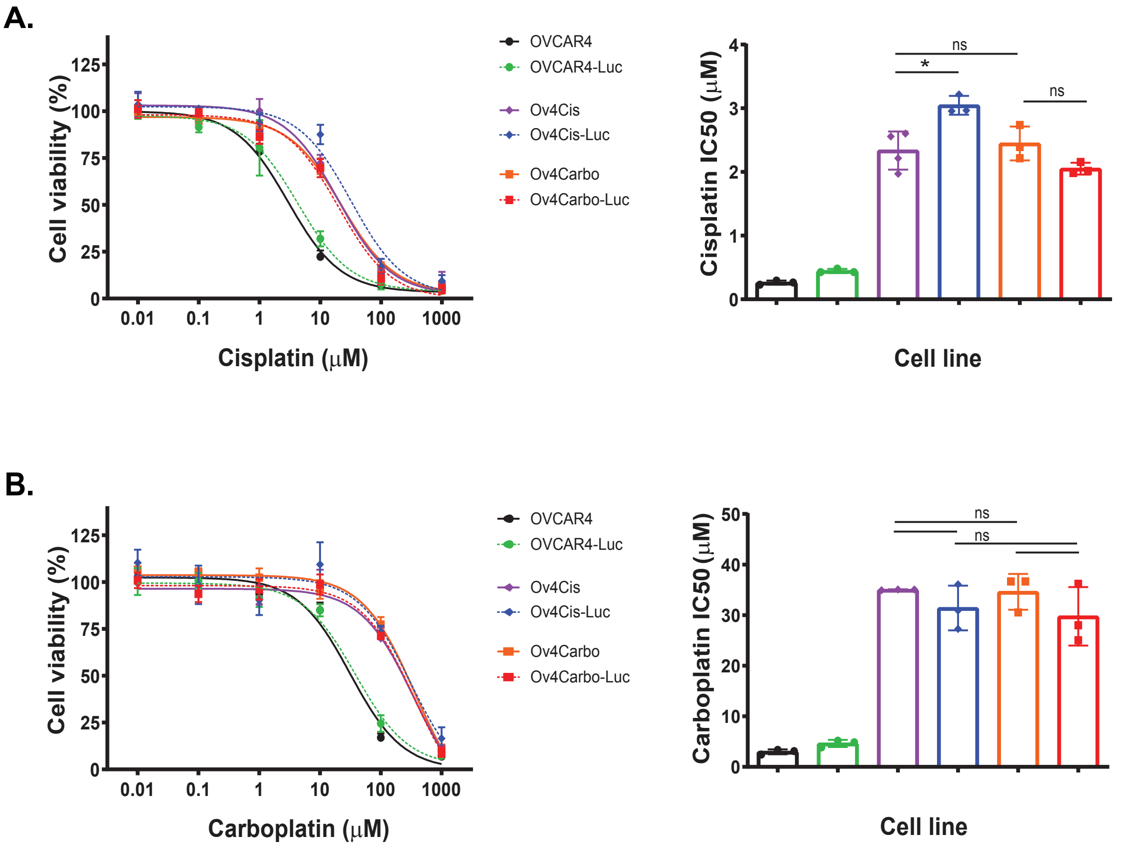

**Supplemental Figure 4**

**A.**

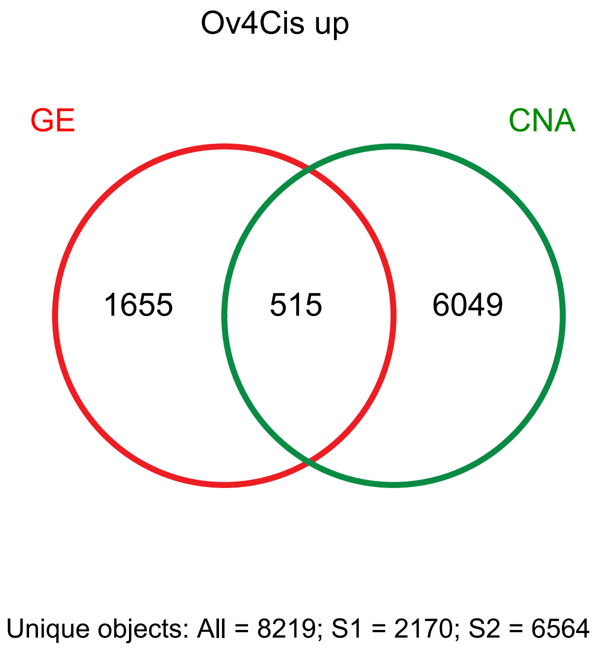


**B.**

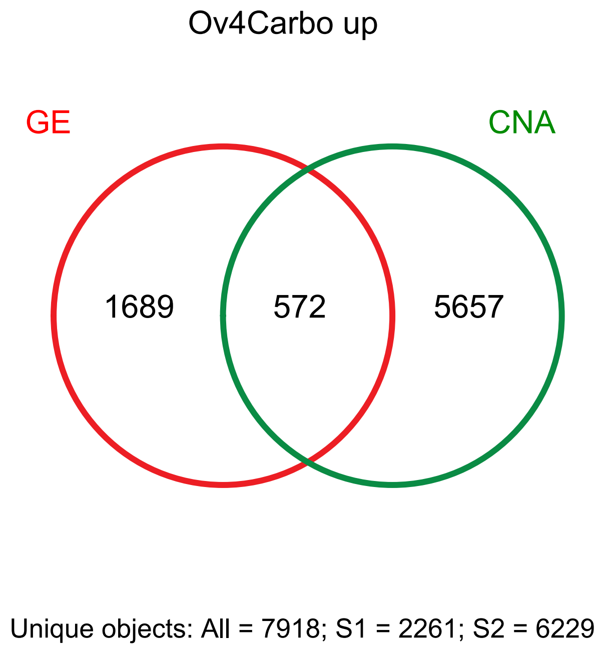


**C.**

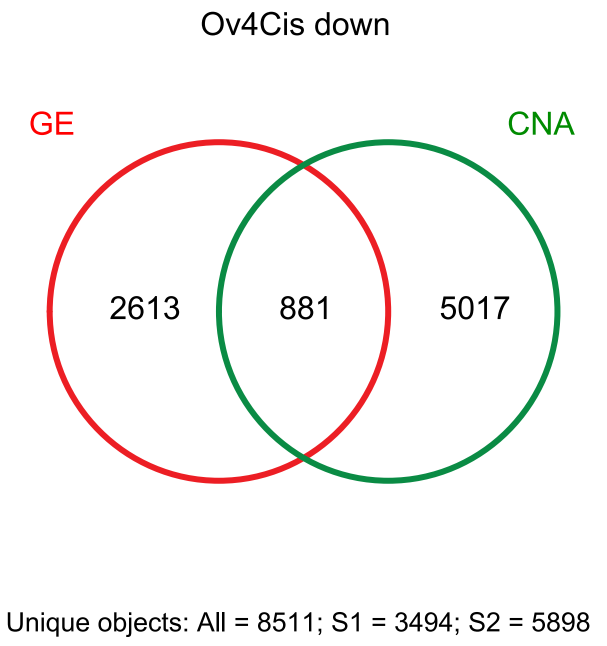


**D.**

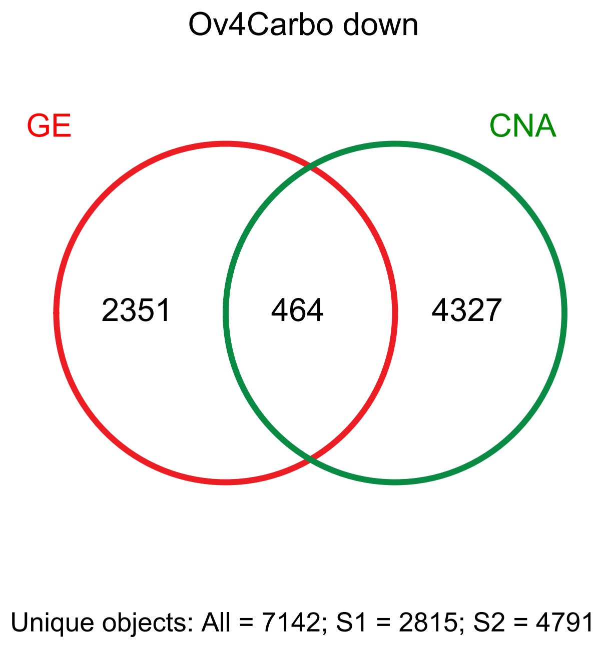

**Supplemental Figure 1:** Cell spheroids were created by growing cells in ultra-low attachment flasks in media supplemented with 1% insulin-transferrin-glucose, 2% B-27, 20ng/ml EGF and 20ng/ml FGF and imaged every four days with light microscopy. D=day

**Supplemental Figure 2:** Unsupervised clustering of cell lines. RNASeq was performed on OVCAR4, Ov4Cis and Ov4Carbo as well as two cell lines derived from single cell Ov4Carbo clones (Ov4Carbo4 and Ov4Carbo7). **A**. Hierarchical cluster analysis and **B.** Principal Component Analysis of gene expression across all samples is illustrated.

**Supplemental Figure 3:** Dose response curves and $IC_{50}$ to cisplatin and carboplatin 72 hours after drug administration. Black = OVCAR4, Green = OVCAR4-Luc, purple = Ov4Cis, blue = Ov4Cis-Luc, orange = Ov4Carbo, red = Ov4Carbo-Luc. $N$=3, mean ± s.d., unpaired $t$-test, ns = non-significant, *$P$<0.05.

**Supplemental Figure 4:** Overlap of genes with concordant copy number variation and significant differential gene expression in resistant (Ov4Cis and Ov4Carbo) *vs.* sensitive OVCAR4 cells. S1 = gene expression (red), S2 = copy number variation (green).
