## supplemental table legends for "Platinum resistance induces diverse evolutionary trajectories in high grade serous ovarian cancer"

**Supplementary Table 1:** List of significantly differentially expressed protein-coding genes in carboplatin resistant Ov4Carbo *vs.* sensitive OVCAR4 and in cisplatin resistant Ov4Cis *vs.* sensitive OVCAR4 cells.

**Supplementary Table 2:** Concordantly changing genes and pathways in resistant *vs.* sensitive OVCAR4-derived cell lines and ICGC resistant/refractory *vs.* sensitive human HGSC.

**Supplementary Table 3:** Overlap of genes with non-synonymous mutations and differential gene expression in resistant Ov4Cis *vs.* sensitive OVCAR4 cells.

**Supplementary Table 4:** Overlap of genes with concordant copy number variation and significant differential gene expression in resistant (Ov4Cis and Ov4Carbo) *vs.* sensitive OVCAR4 cells.

**Supplementary Table 5:** Gene Ontology Biological Process (GOBP) enrichment analysis of genes with copy number alterations that were reflected in concordant gene expression differences between resistant and sensitive OVCAR4 cells.

**Supplementary Table 6:** List of genes in heatmap clusters.

**Supplementary Table 7:** List of significantly differentially expressed protein-coding genes in all platinum resistant cells lines *vs*. the sensitive OVACR4 parental cell line.

**Supplementary Table 8:** Significantly up-regulated or down-regulated pathways ($P$<0.05) between platinum resistant cells lines (Ov4Cis and Ov4Carbo) and sensitive OVCAR4 cells.
